# The Genomic Landscape of Centromeres in Cancers

**DOI:** 10.1101/505800

**Authors:** Anjan K. Saha, Mohamad Mourad, Mark H. Kaplan, Ilana Chefetz, Sami N. Malek, Ronald Buckanovich, David M. Markovitz, Rafael Contreras-Galindo

**Author notes:** Correspondence: Rafael Contreras-Galindo, The Hormel Institute, University of Minnesota, Austin, MN, 55912, USA.

## Abstract

Centromere genomics remain poorly characterized in cancer, due to technologic limitations in sequencing and bioinformatics methodologies that make high-resolution delineation of centromeric loci difficult to achieve. We here leverage a highly specific and targeted rapid PCR methodology to quantitatively assess the genomic landscape of centromeres in cancer cell lines and primary tissue. PCR-based profiling of centromeres revealed widespread heterogeneity of centromeric and pericentromeric sequences in cancer cells and tissues as compared to healthy counterparts. Quantitative reductions in centromeric core and pericentromeric markers (α-satellite units and HERV-K copies) were observed in neoplastic samples as compared to healthy counterparts. Subsequent phylogenetic analysis of a pericentromeric endogenous retrovirus amplified by PCR revealed possible gene conversion events occurring at numerous pericentromeric loci in the setting of malignancy. Our findings collectively represent a more comprehensive evaluation of centromere genetics in the setting of malignancy, providing valuable insight into the evolution and reshuffling of centromeric sequences in cancer development and progression.

## INTRODUCTION

The centromere is essential to eukaryotic biology due to its critical role in genome inheritance^1,2^. The nucleic acid sequences that dominate the human centromeric landscape are α-satellites, arrays of ~171 base-pair monomeric units arranged into higher-order arrays throughout the centromere of each chromosome^3–1031–3^. These α-satellites underlie a hierarchical network of proteins that collectively make up the kinetochore, a large multimeric structure that serves as a molecular bridge between chromosomes and microtubule polymers from the mitotic spindle during cell division. The interaction between centromeres, kinetochores and microtubule polymers lies at the nexus of metaphase and anaphase, ensuring faithful separation of the sister chromatids during mitosis.

Centromeres are thus critical to maintaining the fidelity of chromosomal segregation in proliferating tissues. While much is known about the hierarchical network of proteins that epigenetically compartmentalizes centromeres, the genomic foundation of the centromere remains largely uncharted. Centromeres remain a genetic black box that encompasses 2-5% of the human genome^4^. Despite advancements in next-generation sequencing (NGS) technologies, full assemblies of centromeric loci are still unavailable within the latest builds of the human genome, with the exception of a linear assembly of the centromere of chromosome Y^5^. Low complexity genomic regions, characterized by the contiguous arrangement of repetitive sequences, present computational challenges owing to nonunique alignments that are impractical for current informatics pipelines to navigate. Low complexity regions like centromeric loci are consequently excluded from most downstream bioinformatics analyses.

Methodologies that can add resolution to the genomic landscape of the centromere will thus play an integral role in developing a more nuanced understanding of its contribution to health and disease. Recent efforts at overcoming the technical shortcomings of NGS approaches have focused on more conventional molecular biology techniques, including extended chromatin fiber analysis, fluorescent *in-situ* hybridization (FISH), Southern blotting, and polymerase chain-reaction (PCR) based approaches^4,6–13^. Chromatin fiber analysis, FISH, and Southern blotting, while effective for qualitatively and quantitatively characterizing localization and size of given centromeric proteins and sequences, are labor, resource, and time intensive. PCR-based approaches offer expedited evaluation of the centromeric content within any given sample, making it more scalable than chromatin fiber analysis and hybridization-based approaches when evaluating samples derived from human cell lines and tissue. Corroboration of the specificity and sensitivity of PCR-approaches by a number of orthogonal methodologies suggests that using rapid centromere targeted PCR methodologies is a viable strategy for studying centromere genetics^8,9,14,15^.

Applying scalable PCR-based approaches to the assessment of centromere size and structure in different biological settings is therefore critical to contextualizing our knowledgebase on centromere genetics. Diseases of cell division, particularly cancer, remain largely unexplored within the realm of centromere genetics^16–20^. Gaining deeper insight into the contribution of centromere genetics to tumorigenesis and cancer progression thus has the potential to inform novel therapeutic strategies capable of improving long-term outcomes. Unfortunately, the oncogenic potential of centromeric sequences remains undetermined, due to the shortcomings of sequencing methodologies.

Here we report substantial heterogeneity in the centromeric landscape in cancer cell lines and tissues, in terms of copy number differences between tissues as well as differences between cancer cells/tissues and healthy cells. Both solid and hematologic tumors demonstrated marked copy number alterations in centromeric and pericentromeric repeats, as measured by a previously described quantitative centromere-specific PCR assay that targets core centromeric α-satellite DNA as well as pericentromeric human endogenous retrovirus (HERV) DNA^9^. Phylogenetic analysis of HERV sequences in several cancer cell lines suggests that pericentromeric sequences undergo aberrant recombination during tumorigenesis and/or disease progression, consistent with derangements that have been previous reported^12,20–22^. Strikingly, centromeric variation is a feature present across cancer tissue types, including primary tissue samples, providing further substantiation to the notion that genomic instability in centromeres is a ubiquitous occurrence in cancer. Evaluation of the centromeric landscape in the setting of malignancy thus reveals marked genetic alterations that may reflect novel pathophysiologic contributions to the development and progression of cancer.

## RESULTS

### Cancer Cell Lines Demonstrate Heterogeneous Alterations in Centromeric and Pericentromeric DNA

NGS approaches to interrogating genetic alterations in cancer have repeatedly demonstrated ubiquitous genomic instability that is a hallmark of malignancy. However, the lack of an end-to-end assembly of centromeric loci prevents mapping of representative centromeric reads to a standardized reference. We have thus employed a rapid PCR-based approach that we previously described to evaluate the genomic landscape of centromeres and pericentromeres in several human cancer cell lines (Fig. 1). The method was previously validated by comparison to meta-analyses of data from studies using NGS and southern blot, as well as through FISH analysis^9^. The cell lines studied here are representative of a variety of different tissue types, originating from both solid and hematologic malignancies. Our PCR-based methodology unveils significant heterogeneity in the centromeric and pericentromeric content in all 24 chromosomes across tissue types and as compared to healthy cells. This heterogeneity extends to HERVs, such as HERV K111, that we have previously shown to reside in pericentromeric regions. Unsupervised hierarchical clustering of the chromosome specific repeats demonstrates a striking organization to the patterns in centromere heterogeneity, differentiated by the region of the centromere (core or pericentromere) to which each repeat localizes. Similar clustering analysis applied across the different cell lines revealed that heterogeneity in centromeric and pericentromeric content is tissue type agnostic, with the exception of healthy peripheral blood lymphocytes (PBLs) that demonstrate higher relative concordance. The heterogeneity observed reflects a preference for contractions in centromeric and pericentromeric content, consistent across numerous tissue types (Supplementary Fig. S1). More specifically, D13Z1, D10Z1, D2Z1, D3Z1, D8Z2, D16Z2, and K111 demonstrated the most appreciable losses when collectively assessing all tested cancer cell lines. The nomenclature of these α-satellites begins with the letter D, followed by their resident chromosome number (1–22, X or Y), followed by a Z, and a number indicating the order in which the sequence was discovered. Consistent with the global loss of whole chromosomes previously reported in teratocarcinoma cells, we noted widescale loss of centromere arrays in teratocarinoma cell lines derived from male patients in this study (Supplementary Fig. S1)^23–26^. Of note, K111 deletion stood out as ubiquitous across all evaluated cell lines. Collectively comparing normal peripheral blood mononuclear cells (PBMCs) to cancer cell lines, grouped by tissue type, revealed marked reductions in pericentromeric material, using K111 copy number as a surrogate for pericentromeric content (Supplementary Fig. S2)^12,27^.

**Figure 1.**
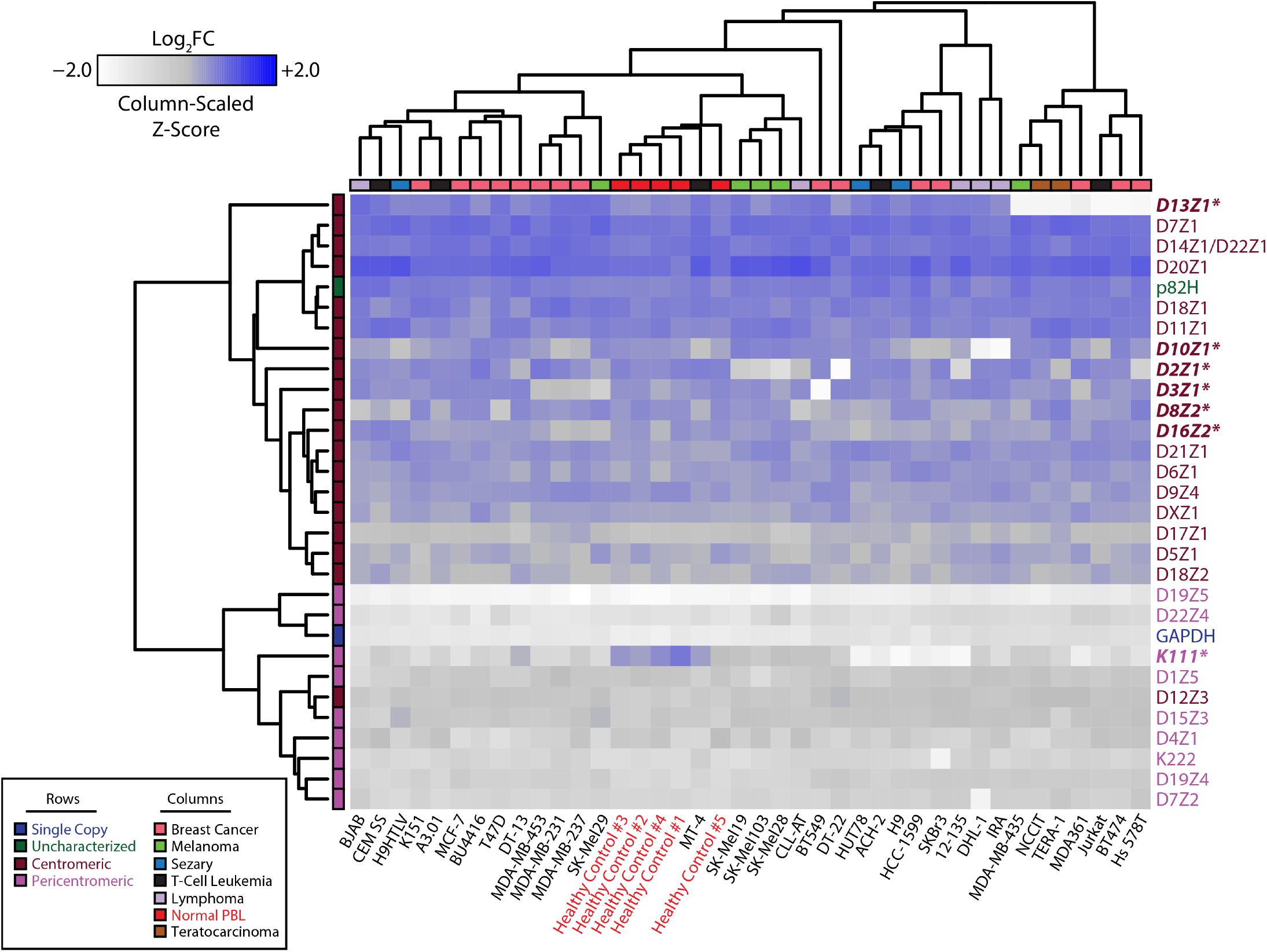
Heterogeneous alterations of centromere DNA in multiple cancer cell lines. Heatmap representing the abundance of α-satellites specific for each centromere array (rows) obtained by qPCR in 50 ng of DNA from healthy cells and from cancer lines (columns). Relative abundance is denoted by the gradient legend (top left). Cancer type and α-satellite localization are depicted as indicated by the legend (bottom left). Repeats marked with an asterisk (also bolded and italicized) represent α-satellites with appreciable alterations across various cell lines relative to healthy controls. Data depicting α-satellite abundance are log_2_ normalized to healthy PBL median values (asterisks, red). The nomenclature of these α-satellites begins with the letter D, followed by their resident chromosome number (1–22, X or Y), followed by a Z, and a number indicating the order in which the sequence was discovered. The DYZ3 repeat was excluded from the analysis to reduce confounding due to gender.

A more focused analysis on breast cancer cell lines allowed us to cross-reference the observed heterogeneity in centromeric DNA against known molecular classifications and karyotypes for each cell line to ascertain whether centromeric and pericentromeric deletions were the result of previously described genetic derangements, such as recurrent molecular alterations or whole chromosome copy number loss, as seen in teratocarcinoma cell lines (Fig. 2)^28–33^. Strikingly, the centromeric content demonstrated heterogeneity across the four molecular subtypes for breast cancer (Basal, HER2, Luminal A, and Luminal B); unsurprisingly, healthy PBLs clustered together. Similar to other tissue types tested, breast cancer cell lines also demonstrated a predilection for contracted centromeres and pericentromeres compared to healthy PBLs (Supplementary Fig. S3). While contraction of D13Z1 in Hs578T, BT474, and MDA-MB-361 can be attributed to loss of whole chromosome 13, contraction of D8Z2 in T47D, D3Z1 and D8Z2 in BT549, and D8Z2 and D10Z1 in SKBr3 were observed despite well characterized copy number amplifications of the respective chromosomes. K111 again demonstrated robust contractions relative to other markers. The strong reduction in DYZ3 (α-satellite on chromosome Y) to nearly undetectable levels provided validation for the specificity of the rapid PCR-based approach to evaluating centromeric content, given the absence of Y-chromosomes in breast cancer cell lines derived from females. Taken together, marked heterogeneity in centromeric and pericentromeric DNA is observed in cancer cell lines, with a predilection towards contraction when comparing cancer cell lines to healthy PBLs.

**Figure 2.**
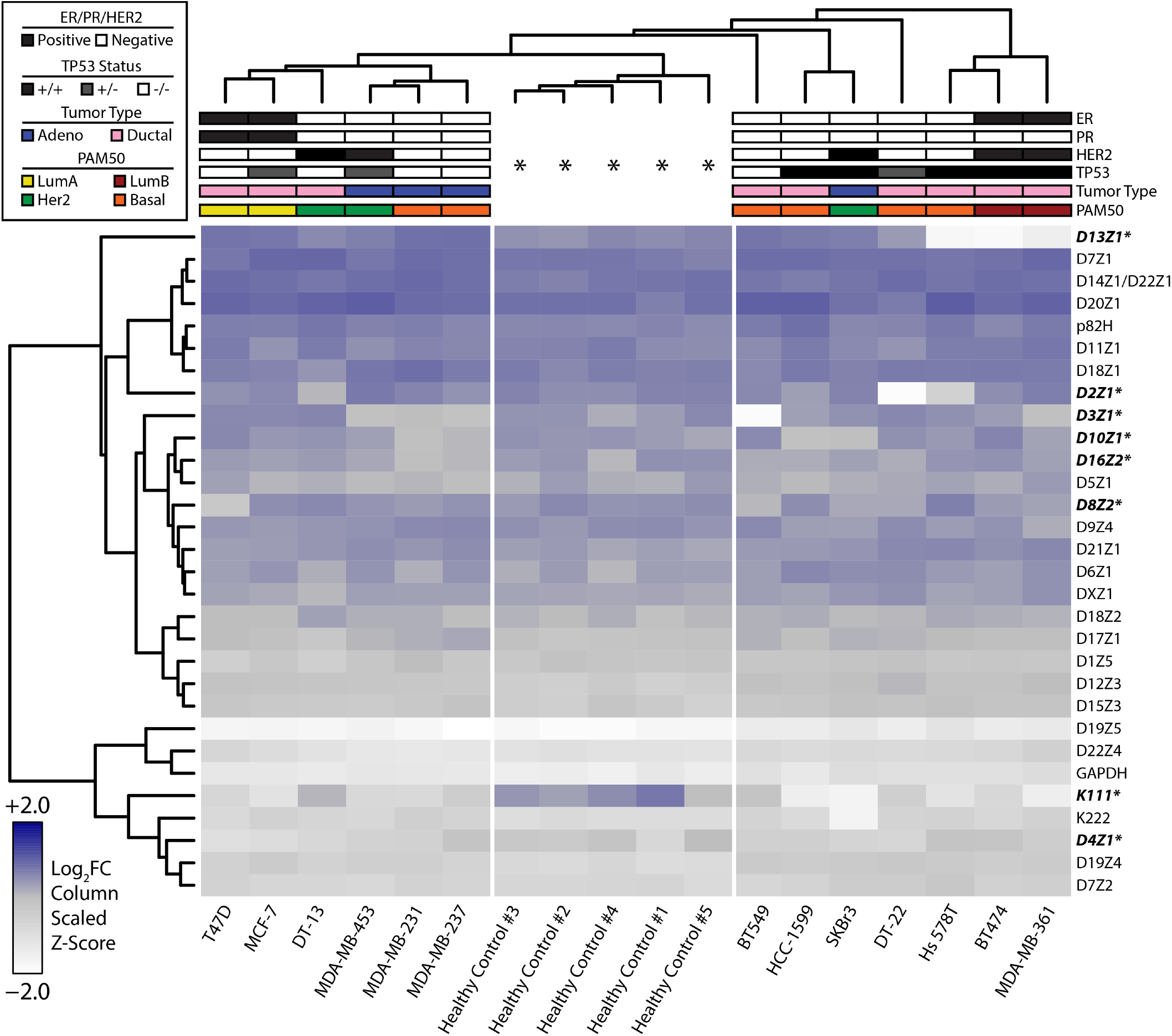
Genomic profiling of centromeres in breast cancer cell lines. Heatmap representing the abundance of α-satellites specific for each centromere array (rows) obtained by qPCR in 50 ng of DNA from healthy cells and from breast cancer lines (columns). Relative abundance is denoted by the gradient legend (bottom left). Data depicting α-satellite abundance are log_2_ normalized to healthy PBL median values (asterisks). Repeats marked with an asterisk (also bolded and italicized) represent α-satellites with appreciable alterations across various cell lines relative to healthy controls. Hormone receptor, *TP53* status, histologic, and molecular classifications are depicted as indicated by the legend (top left). The DYZ3 repeat was excluded from the analysis to reduce confounding due to gender.

### Gene Conversion of Pericentromeric HERV Sequences in Cancer Cell Lines

The genomic landscape of the centromere is characterized by thousands of copies of repetitive elements arranged in tandem to form higher order arrays^1^. Repetitive genomic regions are known to be subject to recombination due to sequence homology^20,34,35^. Intrachromosomal recombination is one example of repeat-associated recombination that can lead to either deletions that reduce the number of repeat units or gene conversion events that genetically homogenize the sequences of repeat units^36–38^. Interestingly, in contrast to healthy PBLs, we identified drastic reductions in pericentromeric K111 sequences across all evaluated cancer cell lines (Fig. 1 and 2). While real-time PCR demonstrates deletion of centromeric and pericentromeric material in cancer cell lines, purely quantitative assessments do not provide insight into other recombination events, such as gene conversion. Furthermore, sequence analysis of α-satellites is unreliable for identifying gene conversion events. We thus conducted phylogenetic analysis on the sequences of real-time PCR amplicons from breast cancer cell lines to identify gene conversion events within K111 loci, given ubiquitous loss of K111 across all cancer cell lines (Fig. 3a). Our previous work has shown that divergence in K111 sequence similarity is dependent on chromosomal location of K111 loci^12,27^. We now show that K111 copies identified in breast cancer cell lines demonstrate cell line dependent sequence convergence towards K111 subtypes that organize into distinct clades (Fig. 3b). The K151 cell line (pink) remarkably produced distinct clades that emerged in close proximity relative to each other from the same ancestral sequence. Sequences amplified from the K151 cell line were notably not distributed heterogeneously throughout the tree. Three additional breast cancer cell lines (MDA-MB-435, DT-13, and HCC1599) formed two exclusive subtypes that were also separated by phylogenetic analysis.

**Figure 3.**
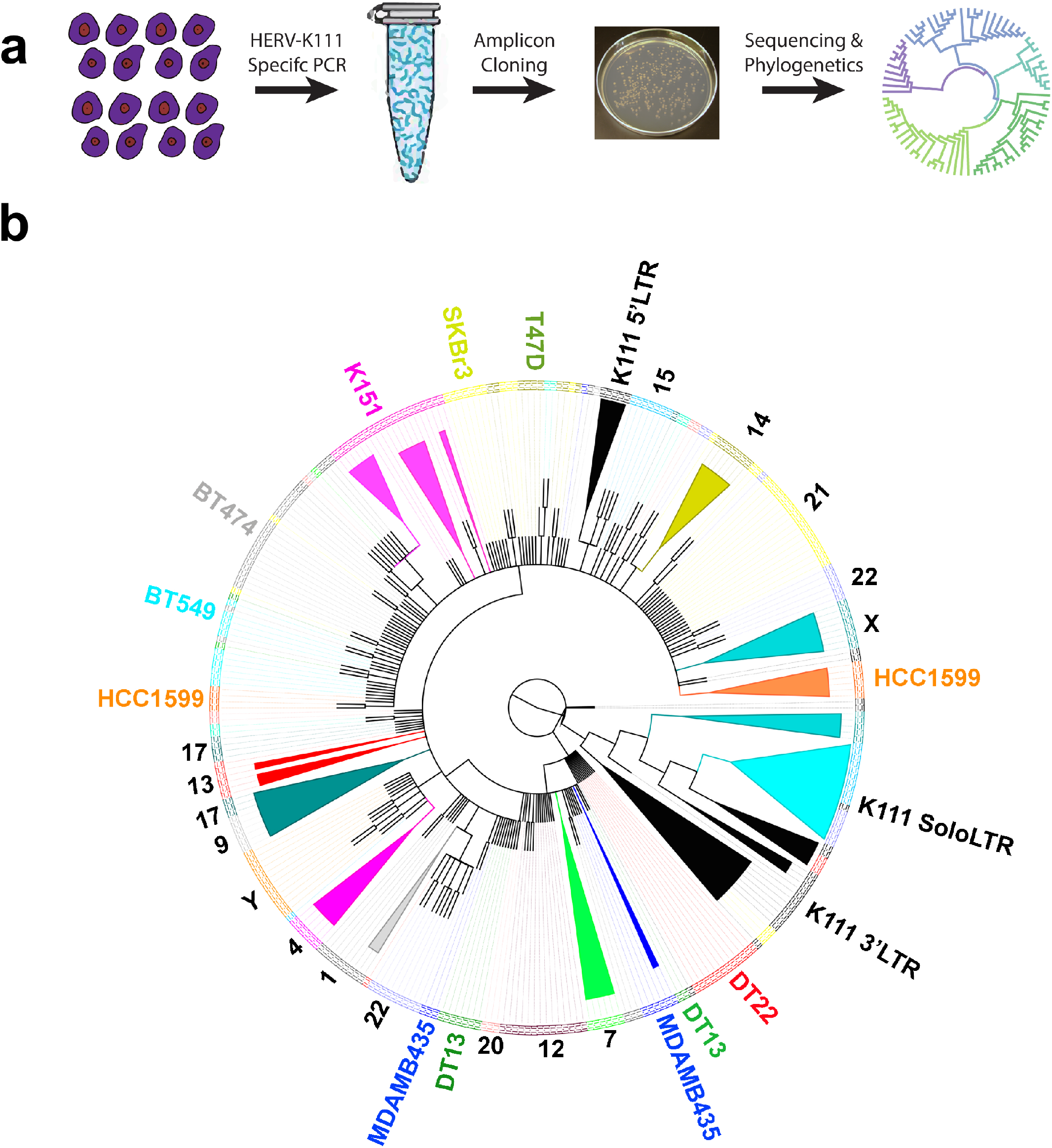
Gene conversion of HERV-K111 in breast cancer cell lines. (**a**) Schematic outline of the experimental methodology employed to identify gene conversion events. (**b**) Phylogenetic analysis conducted on K111 sequences amplified by PCR on breast cancer cell lines (T47D, BT549, HCC-1599, MD-MB-435, DT13, DT22, K151, and SKBr3) and human-hamster hybrid cell lines (each containing a single human chromosome) as a reference. Amplicons are labeled and color-coded along the edge of the phylogenetic tree according to the cell line that produced the amplicon. Amplicons from human-hamster hybrid cell lines are denoted numerically by the human chromosome present in each hybrid cell line. Amplicons from K111 5’LTR, 3’LTR, and Solo LTR are additionally denoted. An example of gene conversion is shown in the cell line K151, possessing clades (pink) that localize in close proximity relative to each other but are not found heterogeneously throughout the tree. Convergence on two distinct K111 subtypes can additionally be identified within the MDA-MB-435, DT-13, and HCC1599 cell lines.

Phylogenetic analysis was also conducted in adult T-cell leukemia (ATL) cell lines and revealed similar patterns as in breast cancer (Supplementary Fig. S4). ATL26 alone formed three exclusive subtypes that diverge in homology from normal K111 clades. Of note, K111 clades arising from ATL43 and ATL16 demonstrated strong homology to K111 Solo LTRs, suggesting intrachromosomal recombination that has deleted K111, i.e. pericentromeric material. ATL43 and ATL16 indeed demonstrate the strongest reductions in K111 copy number relative to other ATL cell lines (Supplementary Fig. S2). As Solo LTRs are the result of homologous recombination between the LTRs flanking endogenous retroviral sequences^39–41^, ATL cell lines having *de novo* K111 sequences with higher relative homology to Solo LTR sequences suggested that pericentromeric K111 sequences served as templates for gene conversion. Taken together, cell line dependent sequence convergence of HERV-K111 in cancer cell lines suggests that gene conversion events are driving sequence evolution within the pericentromeres of cancer cell lines.

### Heterogeneous Loss of Centromere DNA in Cancer Tissue

Human cancer cell lines are useful models for evaluating cancer biology and genetics in an *in vitro* setting. Indefinite cellular propagation, however, results in clonal selection for cells that have a fitness advantage for growing *ex vivo*. Such a fitness advantage is sometimes conferred by abnormal karyotypes (aneuploidy), a cytogenetic feature that can influence the results of PCR based analyses. Cancer tissue itself thus presents the most accurate representation of malignancy-associated genomic instability that results from microenvironmental pressures that cannot be reproduced *ex vivo*. We thus applied our rapid PCR-based approach to DNA isolated from primary cancer tissue. Profiling the centromeric landscape in 9 different ovarian cancer samples against matched PBMCs revealed similarly significant loss of α-satellites across multiple chromosomes as observed in cell lines (Fig. 4). Indeed, quantitative assessment of this heterogeneity again revealed copy number reductions in the cancer tissue, similar to findings noted in cell lines (Supplementary Fig. S5). Strikingly, a drastic reduction in the centromere of chromosome 17 (D17Z1) was seen in ovarian cancer tissue when compared to healthy tissue (Supplementary Fig. S5), corroborating previous reports of chromosome 17 anomalies in ovarian cancer. No changes were seen in the single copy gene *GAPDH* found in the arm of chromosome 12. A significant loss in GAPDH is, however, noted in Sample 285, raising the possibility that this sample’s karyotype displayed derangements that are reflected in the PCR data. Tumor karyotypes for tested samples were, however, unavailable for corroboration.

**Figure 4.**
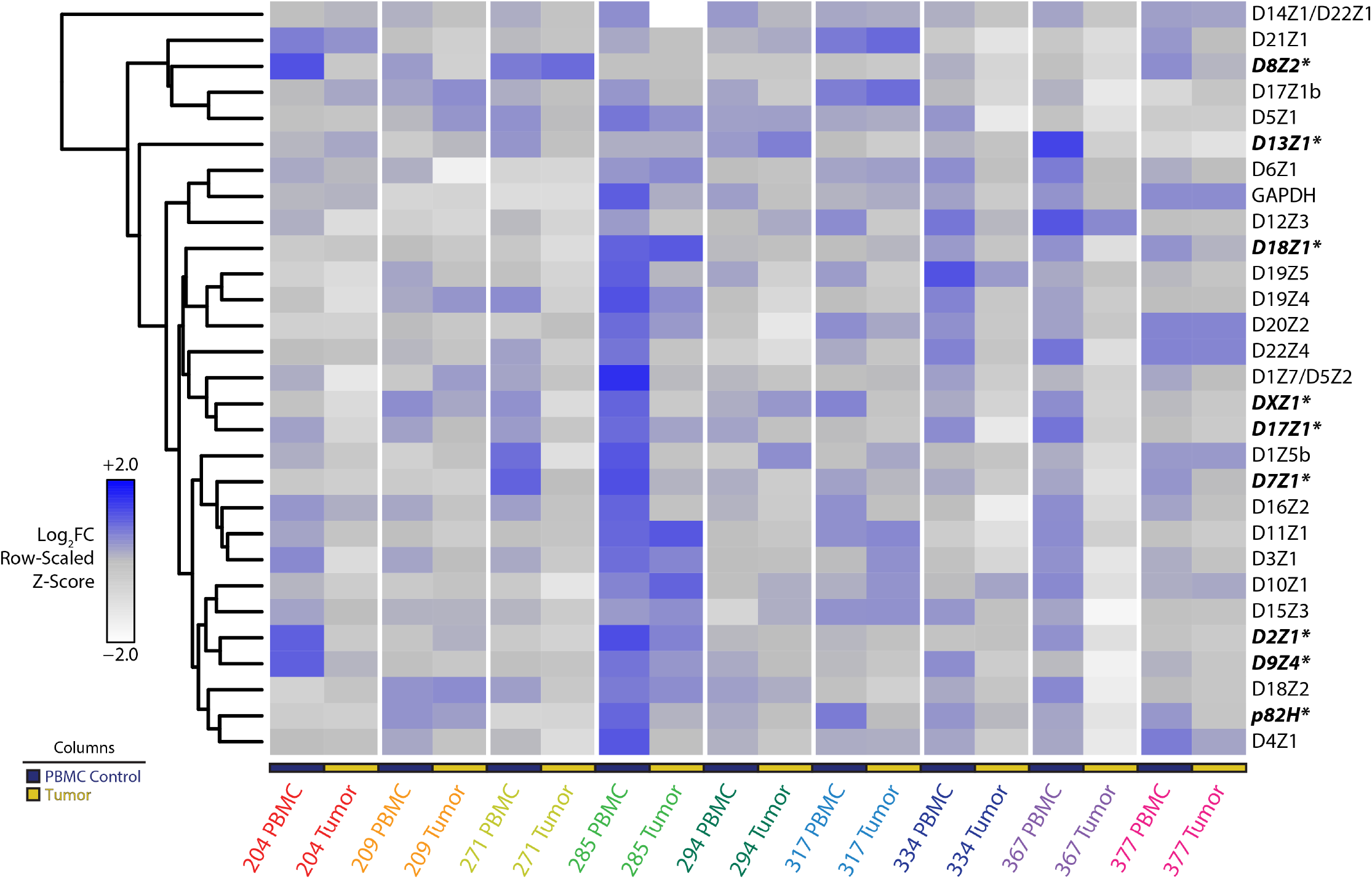
Genomic profiling of centromeres in primary ovarian cancer tissue. Heatmap representation of rapid PCR data from nine primary ovarian cancer tissue samples with matched PBMC DNA. Matched sets from the same patient are grouped by color. PBMC control samples and tumor samples are labeled according to the legend (bottom left). Data depicting α-satellite abundance are log_2_ normalized to PBMC median values. Relative abundance is denoted by the gradient legend (bottom left). Repeats marked with an asterisk (also bolded and italicized) represent α-satellites with appreciable alterations across tissue samples relative to PBMC controls.

While matched blood samples provide reliable non-malignant references to their malignant counterparts, comparisons between primary ovarian cancer tissue and matched blood does not sufficiently deconvolute tissue specific genetic heterogeneity that may be present in normal biologic settings. To expand upon our findings, and to specifically address this latter issue, we profiled the centromeres of B-cells and T-cells that were separated by cell-surface marker selection from chronic lymphocytic leukemia (CLL) primary samples. CLL is a malignancy that arises in B-cells, as opposed to T-cells, within the bone marrow. Applying our methodology to compare patient matched B-cells and T-cells from CLL samples, both cells of lymphocytic lineage, thus largely eliminates the confounding contributions of normal development and tissue specificity to genetic heterogeneity in the centromere. Fewer repeats per sample were evaluated than in the experiments described above due to the limited availability of tumor DNA from each patient. Intriguingly, unsupervised hierarchical cluster analysis across patient samples cleanly separates healthy cells from diseased cells based on chromosome specific α-satellite abundance (Fig. 5). We show contraction of numerous centromeres in malignant CD19+ B-cells as compared to their normal CD3+ T-cell counterparts, whereas no changes were seen in the housekeeping gene *GAPDH* found in the arm of chromosome 12 (Supplementary Fig. S6). Strikingly, we see no such centromeric differences between B-cells and T-cells separated from blood samples derived from healthy individuals. Taken together, centromeric contraction is a characteristic that is present in primary cancer samples, consistent with our data in cancer cell lines.

**Figure 5.**
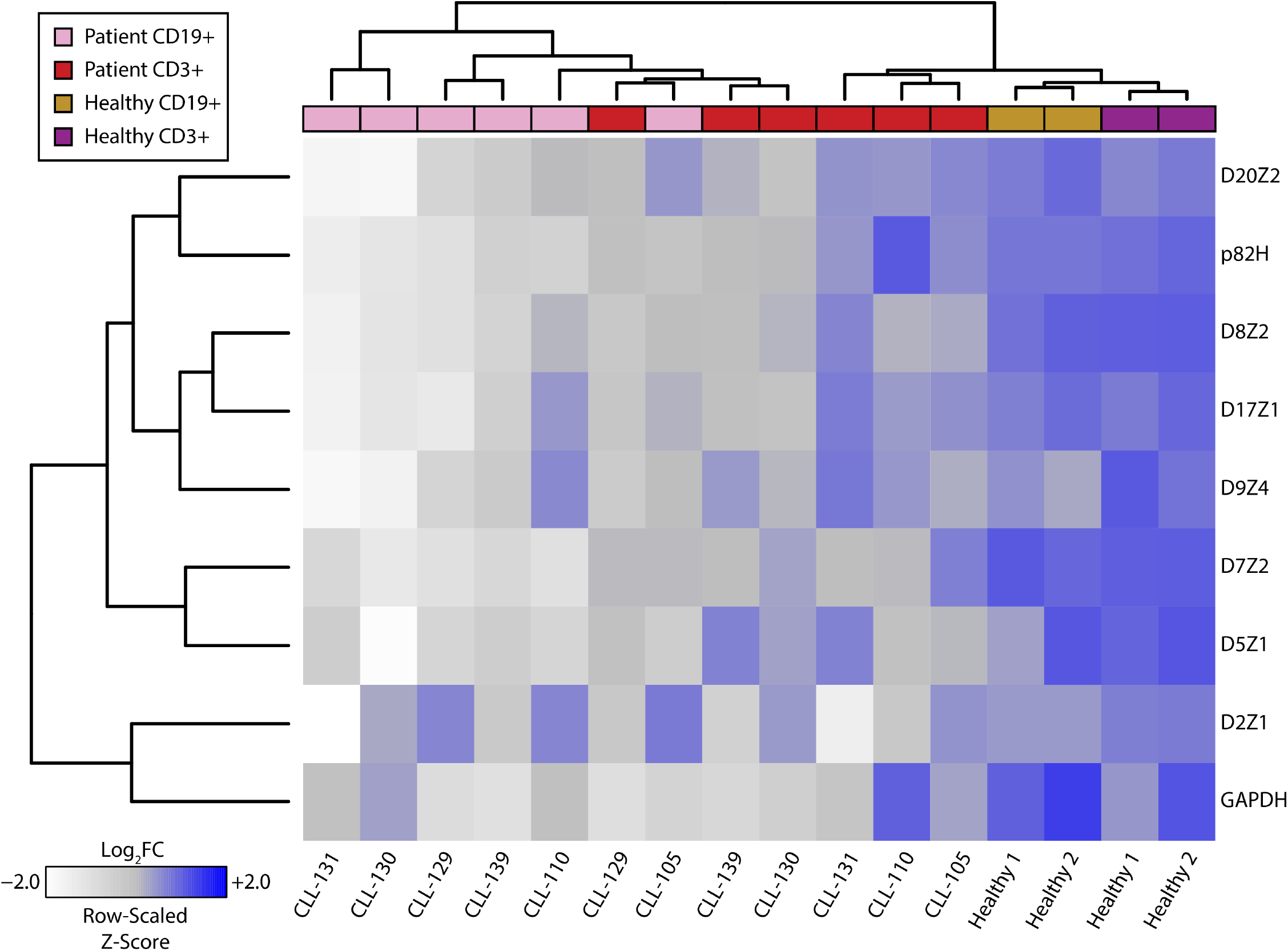
CLL (malignant B-cells) and patient matched T-cells assessed for select centromeric α-satellite markers. Heatmap representation of rapid PCR data from six primary CLL and two healthy samples post-separation by indicated cell surface markers into B-cell (CD19+) and T-cell (CD3+) populations. Data depicting α-satellite abundance are log_2_ normalized to T-cell median values. Relative abundance is denoted by the gradient legend (bottom left). Lymphocyte characterization and disease status is depicted as indicated by the legend (top left).

## DISCUSSION

The importance of centromeres to cell division provides a strong rationale for interrogating the genetics of the centromere in cancers. The challenges associated with studying the genomic landscape of centromeres, owing to the informatics impracticalities of evaluating low complexity regions, have however hindered meaningful progress in understanding the contributions of centromere genetics to tumorigenesis and cancer progression. Only one previous study reported the loss of centromere DNA in leukemia cells using fluorescent in situ hybridization (FISH)^43^. We demonstrate, for the first time, that centromeres and pericentromeres display heterogeneous alterations in the setting of malignancy, both in cancer cell lines and primary samples. We show that these heterogeneous alterations reflect marked reductions and gene conversions of repetitive elements and HERVs in multiple centromeres and pericentromeres, suggesting that oncogenic genomic instability selects against the presence of most centromeric sequences and perhaps for certain pericentromeric sequences. While mechanistically uncharacterized, these findings have direct implications for our understanding of global genomic instability in cancer, given the importance of centromeres to faithful segregation of chromosomes. The loss of centromeric material in chromosome 17 described above is an example of the concordance between centromere instability and ovarian cancer pathogenesis, given the recurrent alterations in chromosome 17 that have been previously described in ovarian cancer^42^. While in some cases loss of centromeric DNA could be attributed to a loss of that entire chromosome, there is also a substantial loss of centromeric DNA in specific chromosomes that are known to be euploid or even polyploid in a given cancer cell line. Further, we have shown previously that DNA from patients with trisomy 13 and trisomy 21 exhibit loss of pericentromeric K111 and that DNA from patients with trisomy 21 exhibit loss of D21Z1, suggesting that pericentromeric and centromeric contraction may drive mis-segregation of chromosomes 13 and 21^9^. It is thus conceivable that alterations in centromeres and pericentromeres may underlie chromosome segregation defects that are routinely observed in the context of abnormal cell proliferation. Gaining deeper insight into the mechanism driving gene conversion and centromere contraction may facilitate the identification of novel molecular drivers that can be targeted to prevent potentially oncogenic mis-segregation events.

While the genetics of centromeres in cancer continue to be elucidated, there is a body of work that has uncovered dysregulation of centromere epigenetics and transcriptional activity in malignancy. Overexpression of CENPA is observed ubiquitously across various cancers, with evidence of ectopic CENPA deposition at extra-centromeric loci across the human genome^44–47^. Satellite RNA abundance is an additional feature that been identified in cell lines and tissue^48–50^. Our findings of genomic contraction of centromeres provides a topographic rationale for the redirection of unbound CENPA to readily accessible ectopic loci in the setting of CENPA overexpression, though additional work is required to distinguish the role of cancer specific post-translational modifications in ectopic deposition of CENPA^51,52^. Moreover, while not mechanistically validated, regions that repress transcriptional homeostasis within centromeric loci may be lost (but beyond the sensitivity of PCR interrogation) during genomic contraction of centromeres and pericentromeres in cancer, thus driving transcriptional activity and overexpression of satellite RNAs in malignancy. Indeed, DNA methylation, an epigenetic mark of transcriptional repression, is prevalent within centromeric loci^53,54^. Selective deletion of methylated regions in centromeres during cancer pathogenesis may relieve transcriptional repression, resulting in overexpression of satellite RNAs. Cancer specific examination of DNA-methylation at the centromeric region that leverages our PCR methodology will be essential to validating this line of reasoning.

Instability in centromeric and pericentromeric loci in the setting of malignancy is consistent with the global genome instability that is a well characterized hallmark of cancer^55^. Subsets of breast and ovarian cancer have well studied DNA repair aberrations in homologous recombination proteins BRCA1/2^56^. Recent genomic profiling of several other malignancies has identified new disease subsets classified by molecular alterations in DNA repair genes and pathways^57,58^. It is conceivable that subsets of cancer that are dysfunctional in DNA repair may exhibit pronounced heterogeneity in centromeric content. Thus, it must be acknowledged that hypermutability in DNA that results from DNA repair dysfunction in cancer may alter centromere and pericentromere sequences enough to prevent detection by PCR, appearing like copy number loss or gene conversion in phylogenetic analysis instead of mismatches or single nucleotide polymorphisms (SNPs). Stratifying samples by DNA repair signatures prior to profiling the genomic landscape of centromeres may provide a strategy for identifying mechanistic contributors to centromere contraction in the setting of malignancy. Moreover, genomic profiling of centromeres in cancer tissue may produce signatures that are predictive of responders to therapies that target the DNA repair machinery, such as poly-ADP ribose polymerase (PARP) inhibitors.

In conclusion, we here provide quantitative resolution of the largely uncharacterized human centromere in the setting of cancer. We notably shed light on a region that has been widely considered a black box and impervious to rapid and comprehensive inquiry at the genomic level. The wide-spread alterations observed in cancer cell lines and primary tissue provide a sound rationale to mechanistically interrogate the molecular machinery that is likely driving the selection against centromeric material. Mechanistic characterization of genomic instability at centromeric loci has the potential to inform therapeutic approaches aimed at improving disease outcomes across several cancer types.

## MATERIALS AND METHODS

### Cell Lines and Cell Culture

Cell lines were cultured according to American Type Culture Collection (ATCC) recommendations. Cell lines were grown at 37 °C in a 5% CO_2_ cell culture incubator and authenticated by short tandem repeat (STR) profiling for genotype validation at the University of Michigan Sequencing Core. ATL cell lines were cultured and authenticated as previously described^59^.

### DNA Isolation

DNA extraction was performed on cell lines and tissue with the DNeasy Blood and Tissue Kit (QIAGEN) according to manufacturer’s instructions. DNA was preserved at −20° C.

### Blood and Tumor Cell Separation

Between January 2005 and September 2016 patients with chronic lymphocytic leukemia (CLL) evaluated at the University of Michigan Comprehensive Cancer Center were enrolled onto this study. The trial was approved by the University of Michigan Institutional Review Board (IRB no. HUM00045507). Patients consented for tissue donation in accordance with a protocol approved by the University of Michigan’s IRB (IRB no. HUM0009149). Written informed consent was obtained from all patients before enrollment in accordance with the Declaration of Helsinki. CLL diagnostic criteria were based on the National Cancer Institute Working Group Guidelines for CLL. Eligible patients needed to have an absolute lymphocytosis (> 5000 mature lymphocytes/μL), and lymphocytes needed to express CD19, CD23, sIg (weak), and CD5 in the absence of other pan-T-cell markers. Peripheral blood mononuclear cells (PBMCs) were isolated by venipuncture and separated using Histopaque-1077 (Sigma). Cryopreserved PBMCs (frozen after Ficoll-gradient purification) from CLL blood specimens were prepared for FACS and sorted into CD19+ (B-cells) and CD3+ (T-cells) cells as previously described^60^. Ovarian cancer DNA were isolated from Stage IIIc or Stage IV ovarian carcinomas. Tumor samples were obtained from the operating room and immediately taken to the laboratory for processing. Tissue was maintained in RPMI/10% FBS throughout processing. Fresh 4 × 4 × 2–mm tumor slices were rinsed several times to remove all loosely attached cells. The tissue was then placed in a tissue culture dish and DNA was extracted as described above.

### Rapid Centromere Target PCR Assay

PCR was conducted on DNA samples from cell lines and primary cancer samples according the previously described conditions^9^. Briefly, copy numbers for each centromeric array, proviruses K111/K222, and single-copy genes were measured by qPCR using specific primers and PCR conditions as described. PCR amplification products were confirmed by sequencing. The qPCR was carried out using the Radiant Green Low-Rox qPCR master mix (Alkali Scientific) with an initial enzyme activation step for 10 min at 95°C and 16–25 cycles consisting of 15 sec of denaturation at 95°C and 30 sec of annealing/extension.

### PCR for 5’ and 3’ K111 LTR Insertions

K111 insertions were amplified by PCR using the Expand Long Range dNTPack PCR kit (Roche Applied Science, Indianapolis, IN) as described. K111 5’ and 3’ LTRs and accompanying flanking regions were amplified. PCR was performed using an initial step of 94 °C for 2 min followed by 35 cycles consisting of denaturation at 94 °C for 30 sec, annealing at 55 °C for 30 sec, and extension at 68 °C for 5 min. The amplification products were cloned into the topo TA vector (Invitrogen, Carlsbad, CA) and sequenced.

### Phylogenetic Analysis

Analysis was conducted as outlined previously^27,61^. The K111-related LTR sequences obtained from the DNA of cell lines, and DNA from human/rodent chromosomal cell hybrids were subjected to BLAST analysis against the NCBI nucleotide database. Sequences were aligned in BioEdit using standard settings and exported to the MEGA5 matrix. LTR trees were generated using Bayesian inference (MrBayes v 3.2^62^) with four independent chains run for at least 1,000,000 generations until sufficient trees were sampled to generate more than 99% credibility. MrBayes integrates the Markov chain Monte Carlo (MCMC) algorithms. MrBayes reads aligned matrices of DNA sequences in standard NEXUS format, so it aligns according to the relative similarity between all the sequences. The trees are unrooted.

### Statistics and Data Analysis

The PCR values obtained in the study were normalized by the total amount of DNA used in the assay as shown in the figure legends. The Z-scores were calculated by determining the number of standard-deviations a copy number value of a given alpha repeat is away from the mean of the values in the same group (cell subset, type of cancer, etc.), assuming a normal distribution. Only the Z-scores in figures S1 to S3 show the standard deviation differences to the mean of the whole data set in order to appreciate the difference in the number of repeats in each centromere array. All heatmaps were generated using the gplots, RColorBrewer, and plotrix packages within the RStudio integrated development environment for the R statistical programming language. Data were log_2_ normalized to the median values of healthy samples. Tests of statistical significance employed two-sided student t-tests, with level of significance denoted on appropriate plots.

### Data Availability

Sequences of K111-related insertions amplified from human DNA and human/rodent somatic chromosomal cell hybrids are deposited in the NCBI database with Accession Numbers (JQ790790 - JQ790967). All other data generated or analyzed during this study are included in this published article (and its Supplementary Information files).

## ACKNOWLEDGEMENTS

The authors are grateful to Debra Buck and Maureen Legendre for assistance and Dr. Gil Omenn for inspiring discussions. ATL cell lines were kindly provided by Dr. Mikulus Popovic from the Institute of Human Virology at the University of Maryland. This work was supported by the National Institutes of Health [K22-CA-177824 to R.C.-G., 5T32GM007863-34, F30-CA-210379 to A.K.S, and RO1-CA-144043 to D.M.M.]; and a University of Michigan Cancer Biology Fellowship [to A.K.S.].

## AUTHOR CONTRUBITIONS

A.K.S. and R.C.-G. conceived, designed and conducted the experiments, analyzed the results, and wrote the main manuscript text; M.M. helped conduct experiments; M.H.K. helped conduct experiments and helped analyze the results; I.C., S.N.M., and R.B. provided patient samples; D.M.M. helped analyze the results and revised the manuscript. All authors reviewed the manuscript.

## REFERENCES

1. Cleveland, D. W., Mao, Y. & Sullivan, K. F. Centromeres and kinetochores: from epigenetics to mitotic checkpoint signaling. Cell 112, 407–421 (2003).

2. Hayden, K. E. Human centromere genomics: now it’s personal. Chromosome Research 20, 621–633 (2012).

3. Henikoff, J. G., Thakur, J., Kasinathan, S. & Henikoff, S. A unique chromatin complex occupies young - satellite arrays of human centromeres. Science Advances 1, e1400234–e1400234 (2015).

4. Aldrup-Macdonald, M. E. & Sullivan, B. A. The past, present, and future of human centromere genomics. Genes (Basel) 5, 33–50 (2014).

5. Jain, M. et al. Linear assembly of a human centromere on the Y chromosome. Nat. Biotechnol. 36, 321–323 (2018).

6. Aldrup-MacDonald, M. E., Kuo, M. E., Sullivan, L. L., Chew, K. & Sullivan, B. A. Genomic variation within alpha satellite DNA influences centromere location on human chromosomes with metastable epialleles. Genome Res. 26, 1301–1311 (2016).

7. Li, X. et al. A fluorescence in situ hybridization (FISH) analysis with centromere-specific DNA probes of chromosomes 3 and 17 in pleomorphic adenomas and adenoid cystic carcinomas. J. Oral Pathol. Med. 24, 398–401 (1995).

8. Liehr, T. Benign and Pathological Chromosomal Imbalances: Microscopic and Submicroscopic Copy Number Variations (CNVs) in Genetics and Counseling. (Academic Press, 2013).

9. Contreras-Galindo, R. et al. Rapid molecular assays to study human centromere genomics. Genome Res. 27, 2040–2049 (2017).

10. Quénet, D. & Dalal, Y. A long non-coding RNA is required for targeting centromeric protein A to the human centromere. eLife 3, (2014).

11. Jabs, E. W., Goble, C. A. & Cutting, G. R. Macromolecular organization of human centromeric regions reveals high-frequency, polymorphic macro DNA repeats. Proc. Natl. Acad. Sci. U.S.A. 86, 202–206 (1989).

12. Zahn, J. et al. Expansion of a novel endogenous retrovirus throughout the pericentromeres of modern humans. Genome Biol. 16, 74 (2015).

13. Du, Y., Topp, C. N. & Dawe, R. K. DNA Binding of Centromere Protein C (CENPC) Is Stabilized by Single-Stranded RNA. PLoS Genetics 6, e1000835 (2010).

14. Shepelev, V. A. et al. Annotation of suprachromosomal families reveals uncommon types of alpha satellite organization in pericentromeric regions of hg38 human genome assembly. Genom Data 5, 139–146 (2015).

15. Miga, K. H. et al. Centromere reference models for human chromosomes X and Y satellite arrays. Genome Res. 24, 697–707 (2014).

16. Vig, B. K., Sternes, K. L. & Paweletz, N. Centromere structure and function in neoplasia. Cancer Genet. Cytogenet. 43, 151–178 (1989).

17. Black, E. M. & Giunta, S. Repetitive Fragile Sites: Centromere Satellite DNA As a Source of Genome Instability in Human Diseases. Genes (Basel) 9, (2018).

18. Bersani, F. et al. Pericentromeric satellite repeat expansions through RNA-derived DNA intermediates in cancer. Proc. Natl. Acad. Sci. U.S.A. 112, 15148–15153 (2015).

19. Natisvili, T. et al. Transcriptional Activation of Pericentromeric Satellite Repeats and Disruption of Centromeric Clustering upon Proteasome Inhibition. PLoS ONE 11, e0165873 (2016).

20. Giunta, S. & Funabiki, H. Integrity of the human centromere DNA repeats is protected by CENP-A, CENP-C, and CENP-T. Proc. Natl. Acad. Sci. U.S.A. 114, 1928–1933 (2017).

21. Yi, J.-M. & Kim, H.-S. Expression and phylogenetic analyses of human endogenous retrovirus HC2 belonging to the HERV-T family in human tissues and cancer cells. J. Hum. Genet. 52, 285–296 (2007).

22. Hughes, J. F. & Coffin, J. M. Human endogenous retroviral elements as indicators of ectopic recombination events in the primate genome. Genetics 171, 1183–1194 (2005).

23. Nathanson, K. L. et al. The Y deletion gr/gr and susceptibility to testicular germ cell tumor. Am. J. Hum. Genet. 77, 1034–1043 (2005).

24. Machiela, M. J. et al. Mosaic chromosome Y loss and testicular germ cell tumor risk. J. Hum. Genet. 62, 637–640 (2017).

25. Mostert, M. M. et al. Fluorescence in situ hybridization-based approaches for detection of 12p overrepresentation, in particular i(12p), in cell lines of human testicular germ cell tumors of adults. Cancer Genet. Cytogenet. 87, 95–102 (1996).

26. Summersgill, B. M. et al. Definition of chromosome aberrations in testicular germ cell tumor cell lines by 24-color karyotyping and complementary molecular cytogenetic analyses. Cancer Genet. Cytogenet. 128, 120–129 (2001).

27. Contreras-Galindo, R. et al. HIV infection reveals widespread expansion of novel centromeric human endogenous retroviruses. Genome Research 23, 1505–1513 (2013).

28. Taherian-Fard, A., Srihari, S. & Ragan, M. A. Breast cancer classification: linking molecular mechanisms to disease prognosis. Brief. Bioinformatics 16, 461–474 (2015).

29. Davidson, J. M. et al. Molecular cytogenetic analysis of breast cancer cell lines. Br. J. Cancer 83, 1309–1317 (2000).

30. Lagos, S. M. R. & Jiménez, N. E. R. Cytogenetic Analysis of Primary Cultures and Cell Lines: Generalities, Applications and Protocols. Recent Trends in Cytogenetic Studies - Methodologies and Applications (2012). doi:10.5772/34200

31. Morris, J. S., Carter, N. P., Ferguson-Smith, M. A. & Edwards, P. A. Cytogenetic analysis of three breast carcinoma cell lines using reverse chromosome painting. Genes Chromosomes Cancer 20, 120–139 (1997).

32. Rummukainen, J. et al. Aberrations of chromosome 8 in 16 breast cancer cell lines by comparative genomic hybridization, fluorescence in situ hybridization, and spectral karyotyping. Cancer Genet. Cytogenet. 126, 1–7 (2001).

33. Letessier, A. et al. Multicolour-banding fluorescence in situ hybridisation (mbanding-FISH) to identify recurrent chromosomal alterations in breast tumour cell lines. Br. J. Cancer 92, 382–388 (2005).

34. Klein, H. L. Genetic control of intrachromosomal recombination. Bioessays 17, 147–159 (1995).

35. Blanco, P. et al. Divergent outcomes of intrachromosomal recombination on the human Y chromosome: male infertility and recurrent polymorphism. J. Med. Genet. 37, 752–758 (2000).

36. Schneider, K. L., Xie, Z., Wolfgruber, T. K. & Presting, G. G. Inbreeding drives maize centromere evolution. Proc. Natl. Acad. Sci. U.S.A. 113, E987–996 (2016).

37. Shi, J. et al. Widespread gene conversion in centromere cores. PLoS Biol. 8, e1000327 (2010).

38. Wolfgruber, T. K. et al. High Quality Maize Centromere 10 Sequence Reveals Evidence of Frequent Recombination Events. Front Plant Sci 7, 308 (2016).

39. Dangel, A. W., Baker, B. J., Mendoza, A. R. & Yu, C. Y. Complement component C4 gene intron 9 as a phylogenetic marker for primates: long terminal repeats of the endogenous retrovirus ERV-K(C4) are a molecular clock of evolution. Immunogenetics 42, 41–52 (1995).

40. Vitte, C. & Panaud, O. Formation of solo-LTRs through unequal homologous recombination counterbalances amplifications of LTR retrotransposons in rice Oryza sativa L. Mol. Biol. Evol. 20, 528–540 (2003).

41. Hughes, J. F. & Coffin, J. M. Human endogenous retrovirus K solo-LTR formation and insertional polymorphisms: implications for human and viral evolution. Proc. Natl. Acad. Sci. U.S.A. 101, 1668–1672 (2004).

42. Tavassoli, M. et al. Whole chromosome 17 loss in ovarian cancer. Genes Chromosomes Cancer 8, 195–198 (1993).

43. MacKinnon, R. N. & Campbell, L. J. The Role of Dicentric Chromosome Formation and Secondary Centromere Deletion in the Evolution of Myeloid Malignancy. Genetics Research International 2011, 1–11 (2011).

44. Sun, X. et al. Elevated expression of the centromere protein-A(CENP-A)-encoding gene as a prognostic and predictive biomarker in human cancers: Elevated Expression of the CENP-A-Encoding Gene in Cancer. International Journal of Cancer 139, 899–907 (2016).

45. Zhang, W. et al. Centromere and kinetochore gene misexpression predicts cancer patient survival and response to radiotherapy and chemotherapy. Nature Communications 7, 12619 (2016).

46. Athwal, R. K. et al. CENP-A nucleosomes localize to transcription factor hotspots and subtelomeric sites in human cancer cells. Epigenetics Chromatin 8, 2 (2015).

47. Lacoste, N. et al. Mislocalization of the centromeric histone variant CenH3/CENP-A in human cells depends on the chaperone DAXX. Mol. Cell 53, 631–644 (2014).

48. Ting, D. T. et al. Aberrant Overexpression of Satellite Repeats in Pancreatic and Other Epithelial Cancers. Science 331, 593–596 (2011).

49. Kishikawa, T. et al. Satellite RNA Increases DNA Damage and Accelerates Tumor Formation in Mouse Models of Pancreatic Cancer. Mol. Cancer Res. 16, 1255–1262 (2018).

50. Kishikawa, T. et al. Satellite RNAs promote pancreatic oncogenic processes via the dysfunction of YBX1. Nat Commun 7, 13006 (2016).

51. Niikura, Y. et al. CENP-A K124 Ubiquitylation Is Required for CENP-A Deposition at the Centromere. Developmental Cell 32, 589–603 (2015).

52. Deyter, G. M. R. & Biggins, S. The FACT complex interacts with the E3 ubiquitin ligase Psh1 to prevent ectopic localization of CENP-A. Genes & Development 28, 1815–1826 (2014).

53. Gopalakrishnan, S., Sullivan, B. A., Trazzi, S., Della Valle, G. & Robertson, K. D. DNMT3B interacts with constitutive centromere protein CENP-C to modulate DNA methylation and the histone code at centromeric regions. Hum. Mol. Genet. 18, 3178–3193 (2009).

54. Kim, I. S. et al. Roles of Mis18α in Epigenetic Regulation of Centromeric Chromatin and CENP-A Loading. Molecular Cell 46, 260–273 (2012).

55. Hanahan, D. & Weinberg, R. A. Hallmarks of cancer: the next generation. Cell 144, 646–674 (2011).

56. Roy, R., Chun, J. & Powell, S. N. BRCA1 and BRCA2: different roles in a common pathway of genome protection. Nat. Rev. Cancer 12, 68–78 (2011).

57. Robinson, D. et al. Integrative Clinical Genomics of Advanced Prostate Cancer. Cell 161, 1215–1228 (2015).

58. Robinson, D. R. et al. Integrative clinical genomics of metastatic cancer. Nature 548, 297–303 (2017).

59. Maeda, M. et al. Origin of human T-lymphotrophic virus I-positive T cell lines in adult T cell leukemia. Analysis of T cell receptor gene rearrangement. Journal of Experimental Medicine 162, 2169–2174 (1985).

60. Kujawski, L. et al. Genomic complexity identifies patients with aggressive chronic lymphocytic leukemia. Blood 112, 1993–2003 (2008).

61. Contreras-Galindo, R. et al. Characterization of human endogenous retroviral elements in the blood of HIV-1-infected individuals. J. Virol. 86, 262–276 (2012).

62. Ronquist, F. & Huelsenbeck, J. P. MrBayes 3: Bayesian phylogenetic inference under mixed models. Bioinformatics 19, 1572–1574 (2003).

