## Supplementary Data for "The Genomic Landscape of Centromeres in Cancers"

### SUPPLEMENTARY FIGURES AND LEGENDS

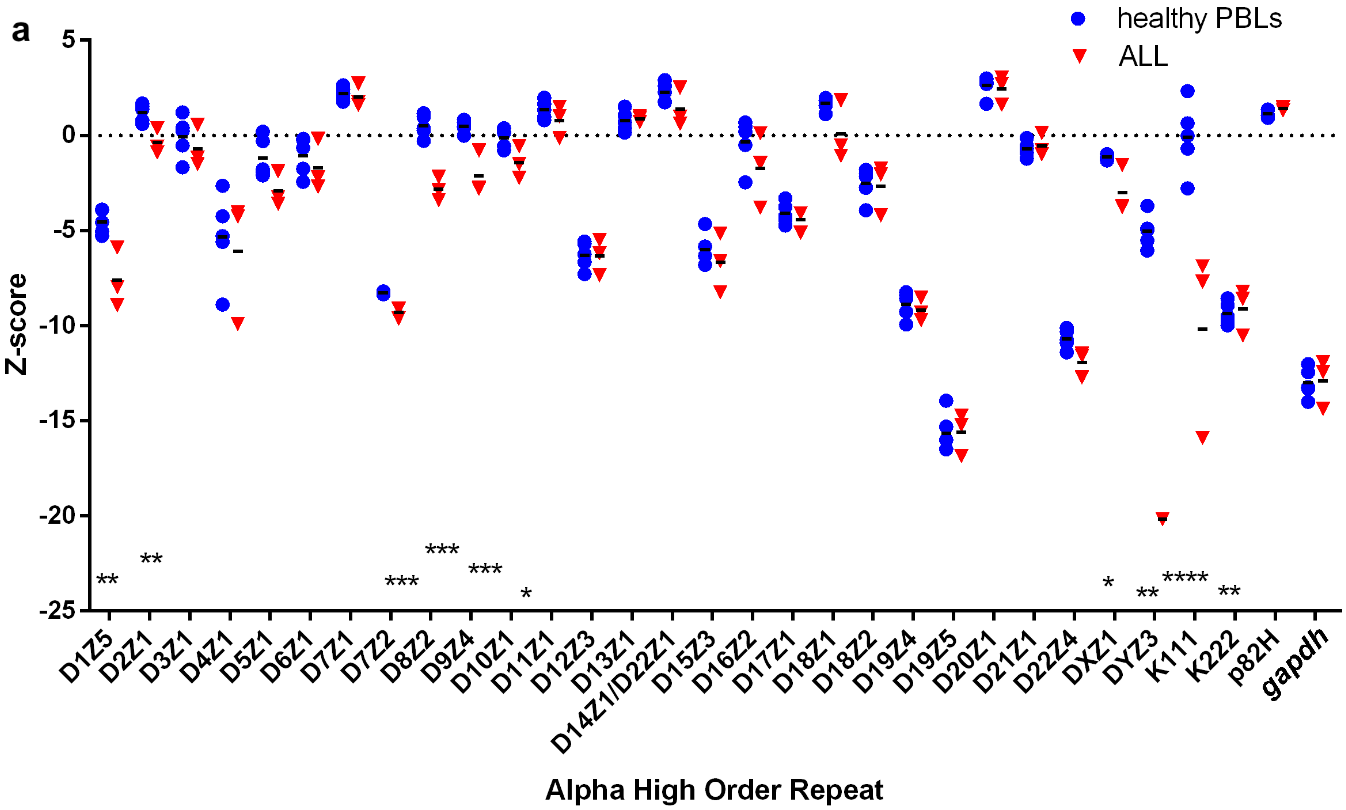

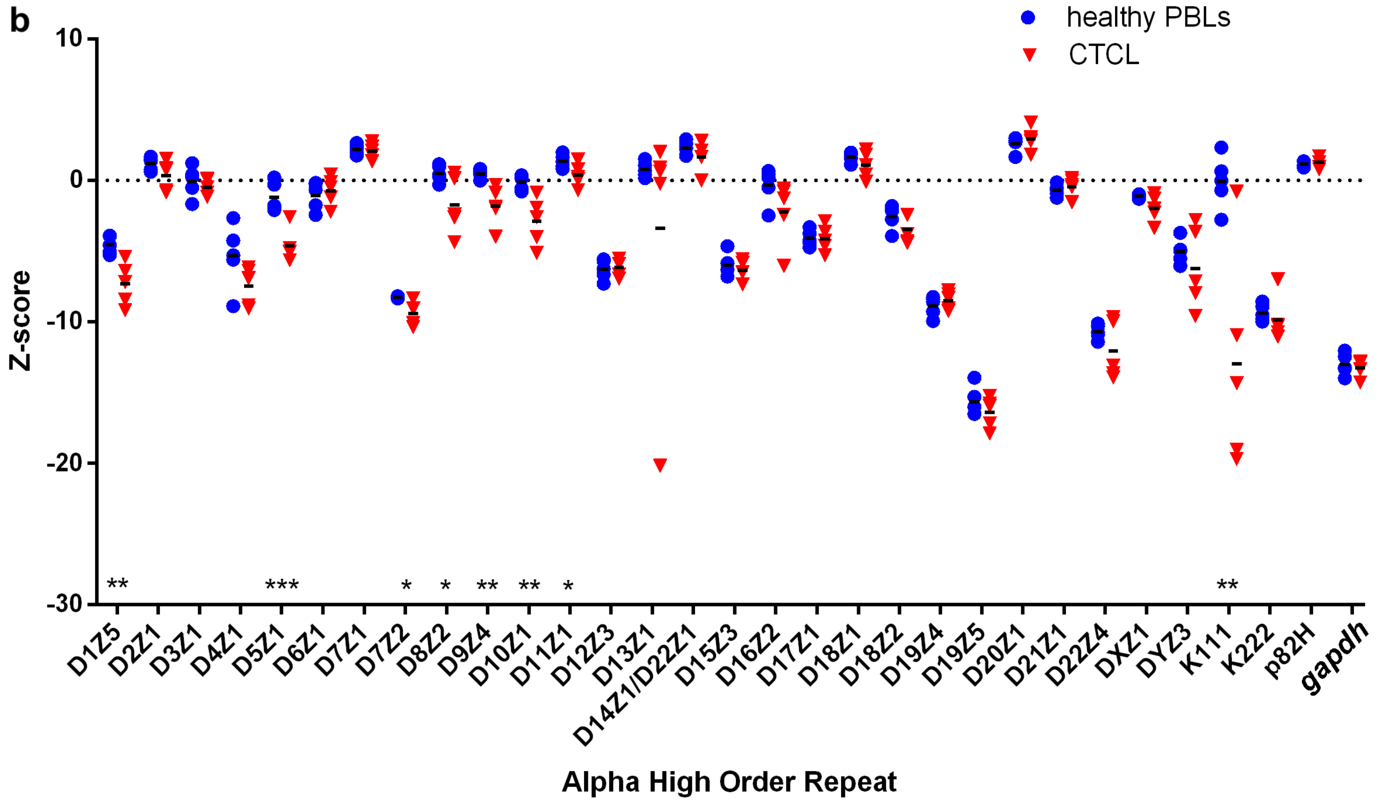
b
healthy PBLs
CTCL
Z-score
10
0
-10
-20
-30
** *** * * ** ** * **
D1Z5 D2Z1 D3Z1 D4Z1 D5Z1 D6Z1 D7Z1 D7Z2 D8Z2 D9Z4 D10Z1 D11Z1 D12Z3 D13Z1 D14Z1/D22Z1 D15Z3 D16Z2 D17Z1 D18Z1 D18Z2 D19Z4 D19Z5 D20Z1 D21Z1 D22Z4 DXZ1 DYZ3 K111 K222 p82H gapdh
Alpha High Order Repeat

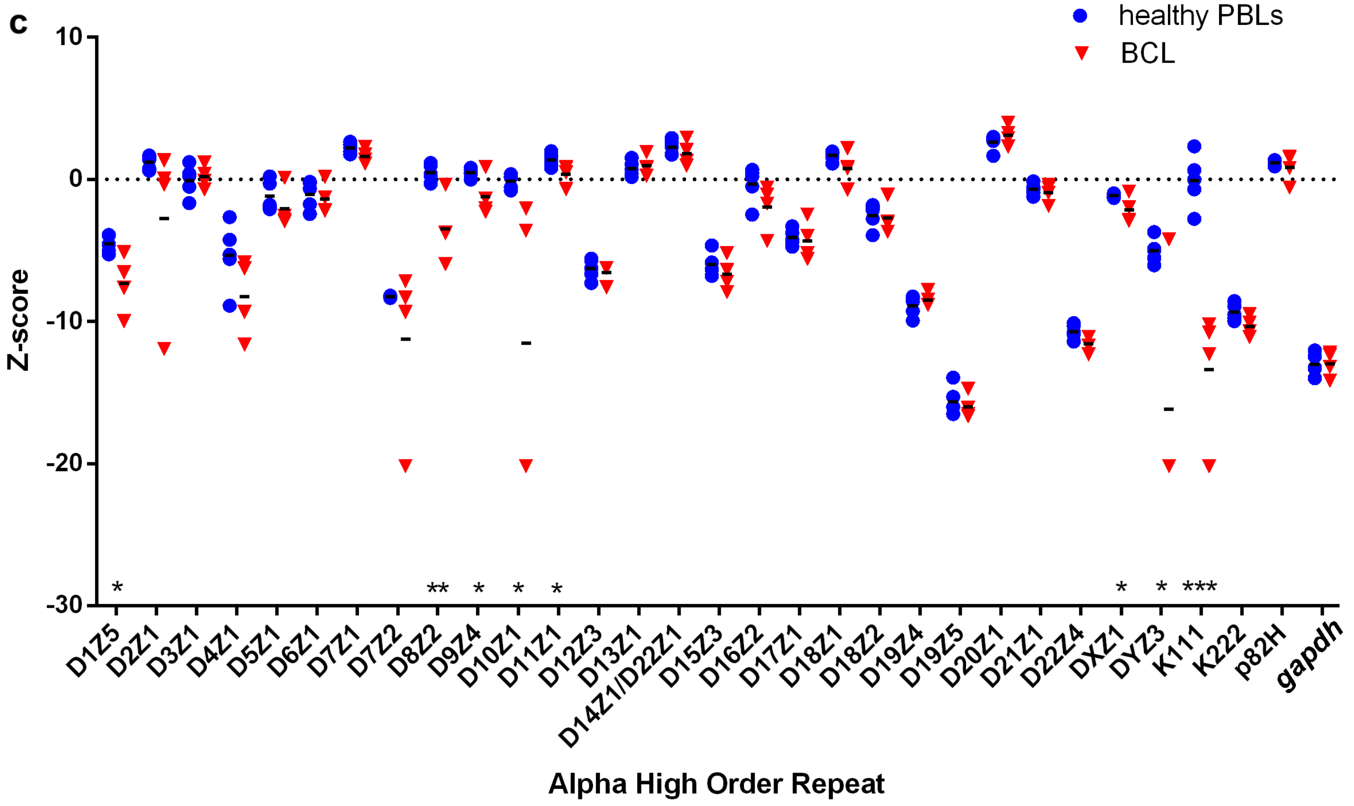
c
healthy PBLs
BCL
Z-score
10
0
-10
-20
-30
* ** * * * * * ***
D1Z5 D2Z1 D3Z1 D4Z1 D5Z1 D6Z1 D7Z1 D7Z2 D8Z2 D9Z4 D10Z1 D11Z1 D12Z3 D13Z1 D14Z1/D22Z1 D15Z3 D16Z2 D17Z1 D18Z1 D18Z2 D19Z4 D19Z5 D20Z1 D21Z1 D22Z4 DXZ1 DYZ3 K111 K222 p82H gapdh
Alpha High Order Repeat

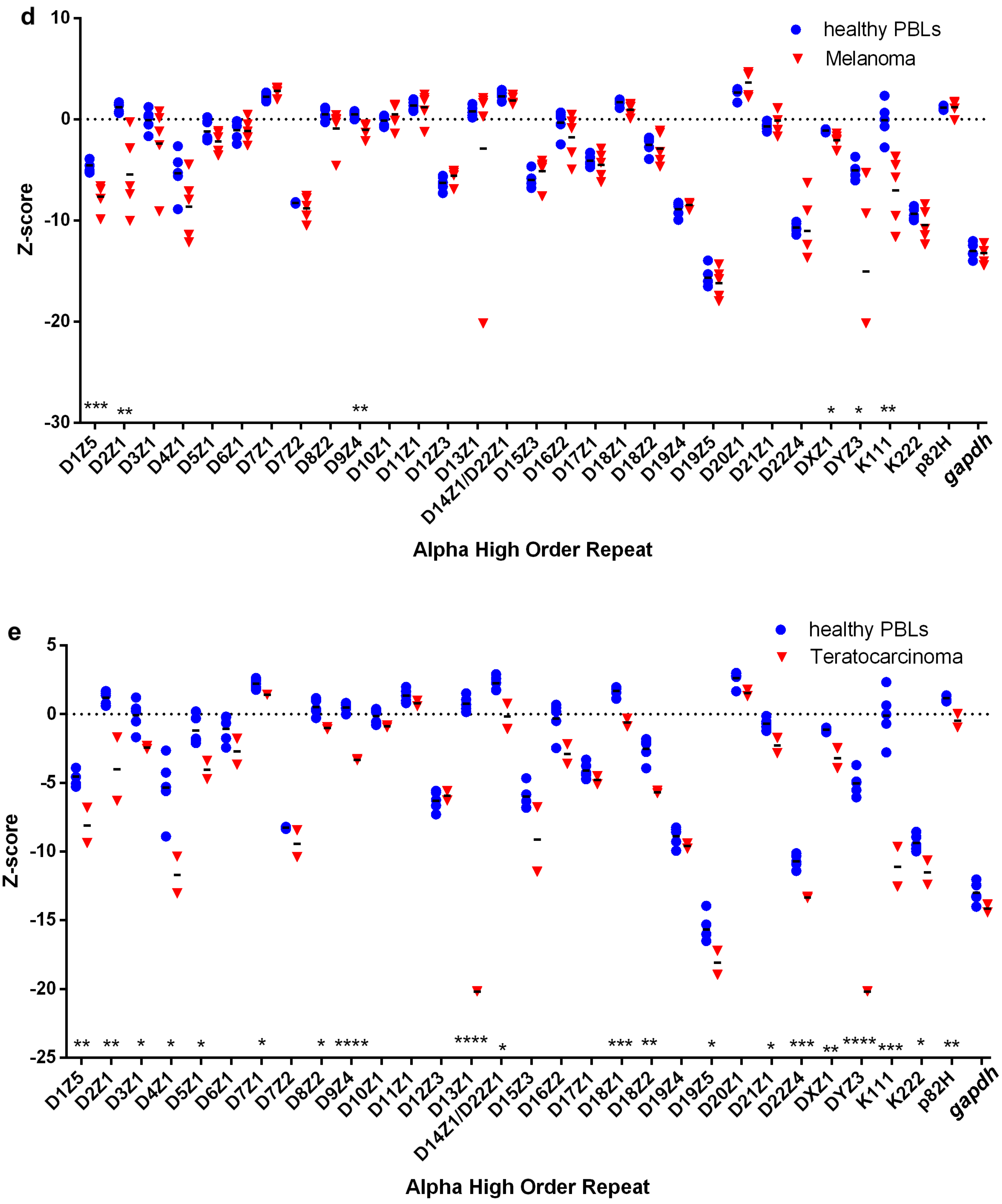


**Supplementary Figure S1.** Heterogeneous loss of centromeres in cancer cell lines. Dot plots representing the abundance of α-satellites specific for each centromere array (X axis) in (**a**) Acute Lymphoblastic Leukemia (ALL) cell lines, (**b**) Cutaneous T-Cell Lymphoma (CTCL) cell lines, (**c**) B-Cell Lymphoma (BCL) cell lines, (**d**) melanoma cell lines, and (**e**) teratocarcinoma cell lines. Abundance of centromere specific α-satellites is depicted by the Z-score (Y-axis) of each α-satellite. The Z-scores show the log number of standard deviation differences to the mean of the whole data set in order to appreciate the difference in the number of repeats in each centromere array. The $\log_2$ normalized numbers for each α-satellite were normalized to the average copy number of repeats in healthy cells (blue circles). Statistical significance was calculated using a t-test. * = $p< 0.05$, ** = $p< 0.01$, *** = $p<0.001$, **** = $p< 0.0001$.

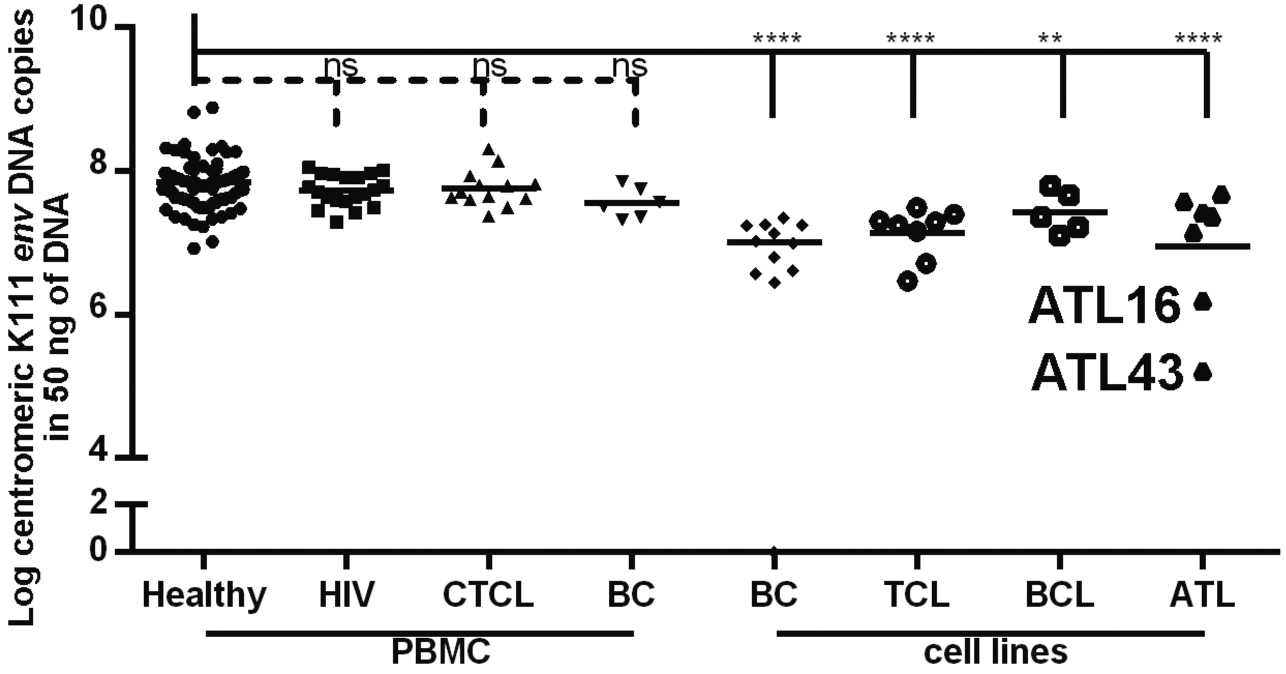


**Supplementary Figure S2.** Abundance of pericentromeric K111 across cell lines and PBMCs. Dot plot representing log-transformed abundance of centromeric HERV *env* sequences from either PBMCs or cell lines with disease type designations listed along the X axis. The *gag* region of K111 was not assessed in our analysis, as select populations and CTCL patients are homozygous null for K111 *gag* (Kaplan et. al. *BMC Medical Genomics*, in press). HIV = HIV patient sample, CTCL = Cutaneous T-Cell Lymphoma, BC = Breast Cancer, TCL = T-Cell Lymphoma, BCL = B-Cell Lymphoma, ATL = Adult T-Cell Lymphoma. Abundance of K111 is depicted by the $log_2$ Z-score (Y-axis). Statistical significance was calculated using a t-test. * = $p< 0.05$, ** = $p< 0.01$, *** = $p<0.001$, **** = $p< 0.0001$.

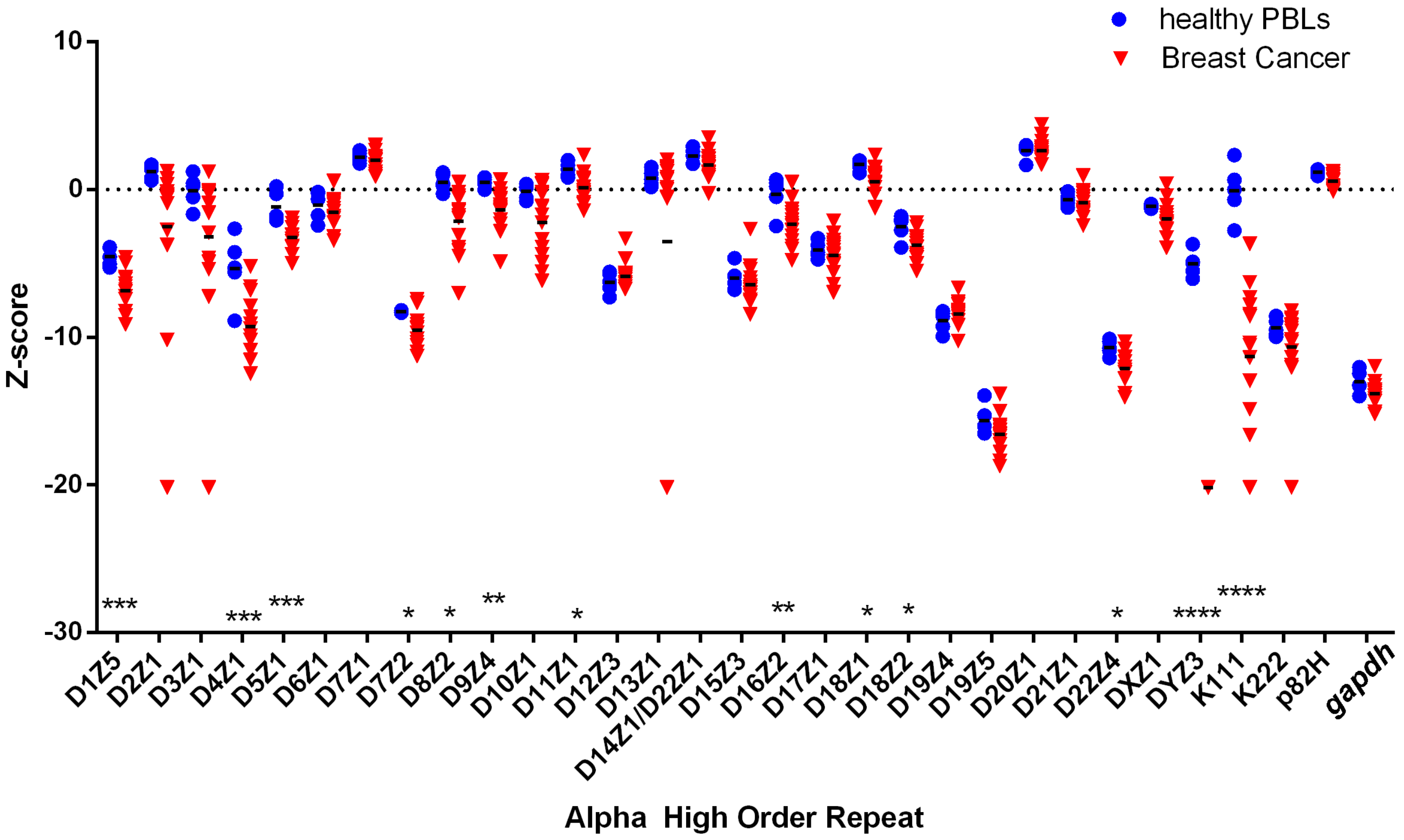


**Supplementary Figure S3.** Heterogeneous loss of centromeres in breast cancer cell lines. Dot plot representing the abundance of α-satellites specific for each centromere array (X axis) in breast cancer cell lines. Abundance of centromere specific α-satellites is depicted by the Z-score (Y-axis) of each α-satellite. The Z-scores show the log number of standard deviation differences to the mean of the whole data set in order to appreciate the difference in the number of repeats in each centromere array. The $\log_2$ normalized numbers for each α-satellite were normalized to the average copy number of repeats in healthy cells (blue circles). Statistical significance was calculated using a t-test. * = $p < 0.05$, ** = $p < 0.01$, *** = $p < 0.001$, **** = $p < 0.0001$.

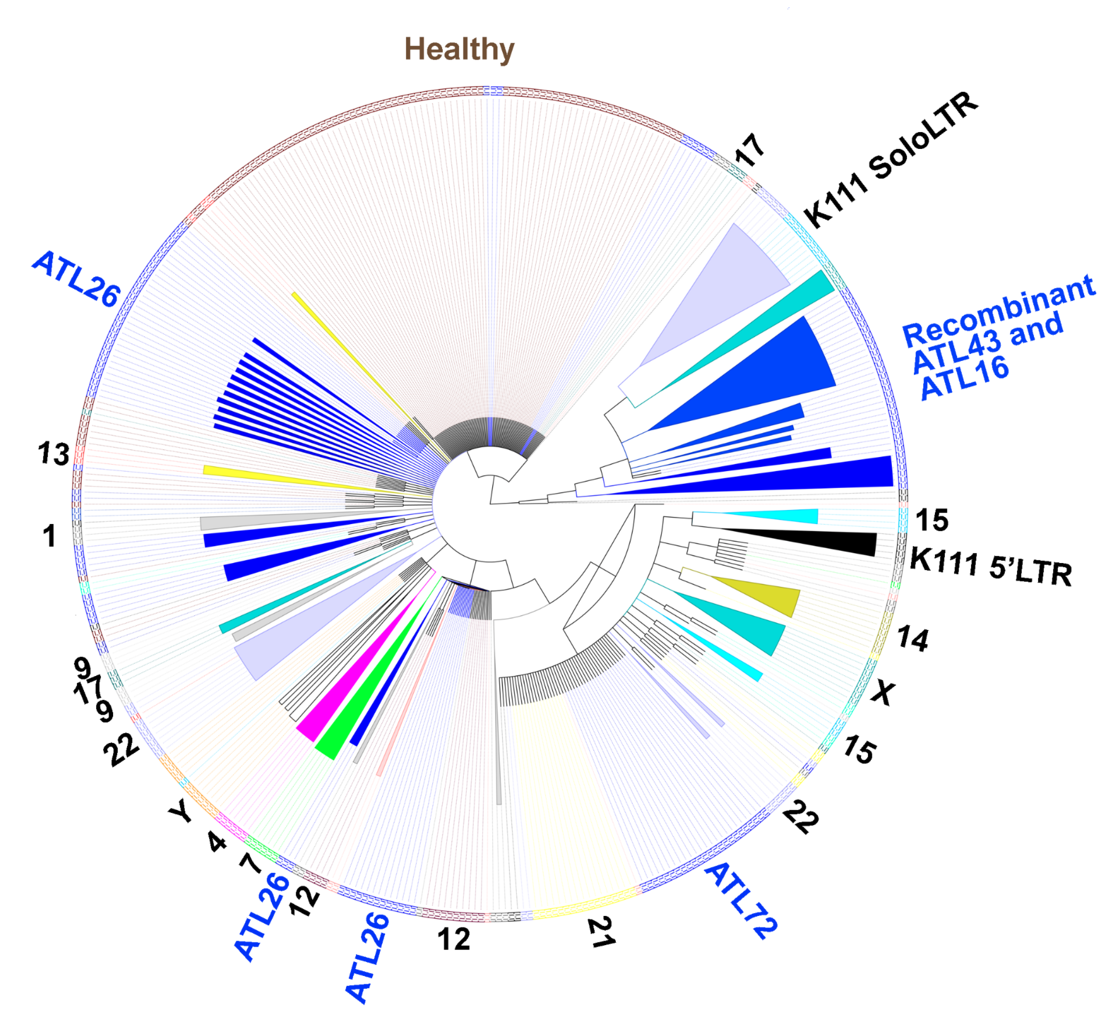


**Supplementary Figure S4.** Gene conversion of HERV K111 in adult T-cell leukemia (ATL) cell lines. Phylogenetic analysis conducted on K111 sequences amplified by PCR in ATL cell lines (ATL26, ATL72, ATL16, and ATL43), human-hamster hybrid cell lines, and heathy cells (brown labels). Amplicons are labeled and color-coded along the edge of the phylogenetic tree according to the cell line that produced the amplicon. Amplicons from human-hamster hybrid cell lines are denoted numerically by the human chromosome present in each hybrid cell line. Amplicons from K111 5’LTR and Solo LTR are additionally denoted. Recombinant K111 sequences resembling Solo LTRs are seen in cell lines ATL43 and ATL16, shown in blue arising from the same ancestral sequence to K111 Solo LTR sequences. K111 sequences from heathy PBLs show heterogeneous distribution along the tree and did not cluster in novel clades.

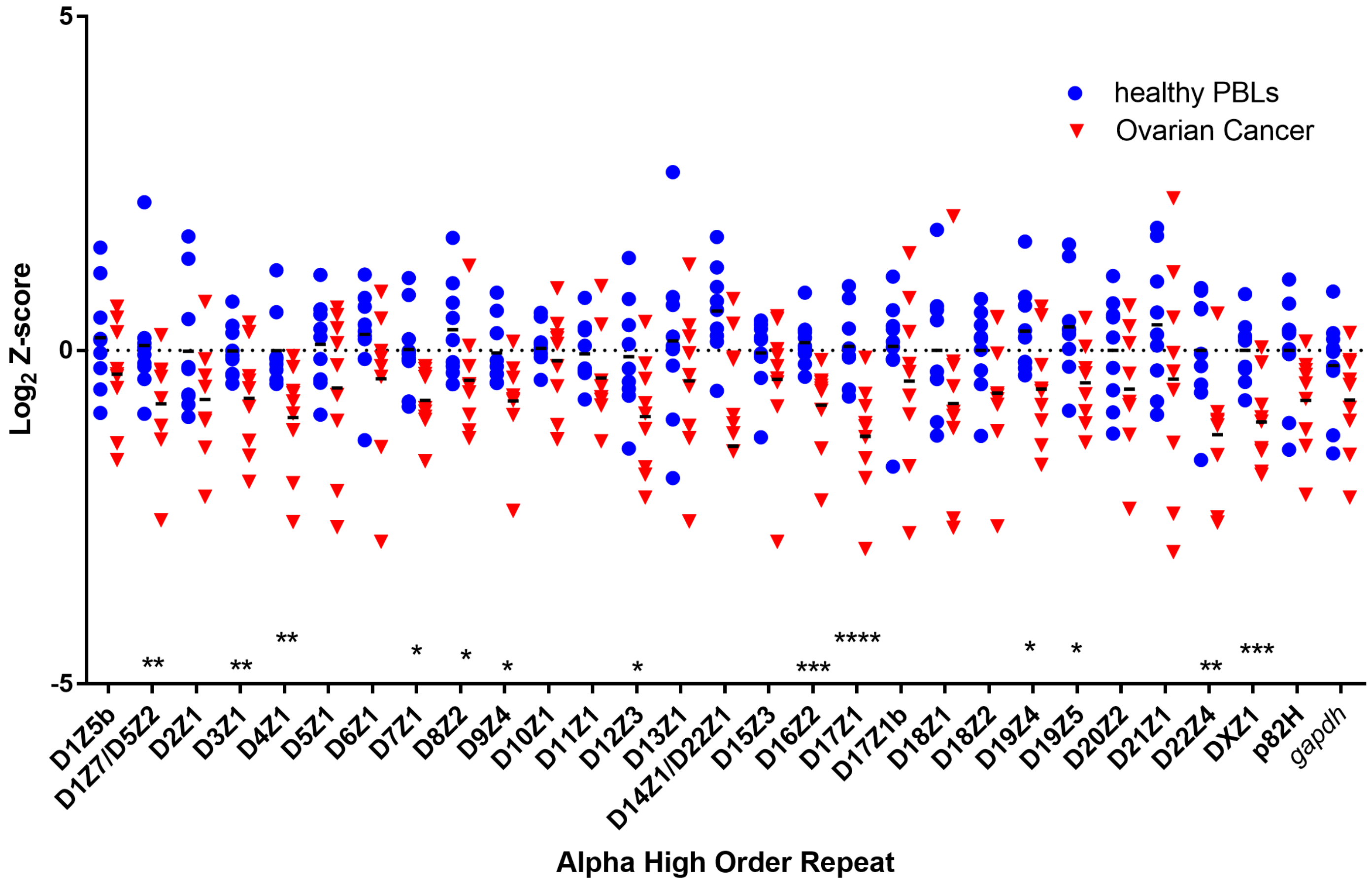


**Supplementary Figure S5.** Heterogeneous loss of centromeres in ovarian cancer patients. Dot plot representing the abundance of α-satellites specific for each centromere array (X axis) in ovarian cancer tumors. Abundance of centromere specific α-satellites is depicted by the $\log_2$ Z-score (Y-axis) of each α-satellite. The $\log_2$ normalized numbers for each α-satellite were normalized to the average copy number of a given repeat in DNA from healthy cells (blue circles). Statistical significance was calculated using a t-test. * = $p$< 0.05, ** = $p$< 0.01, *** = $p$<0.001, **** = $p$< 0.0001.

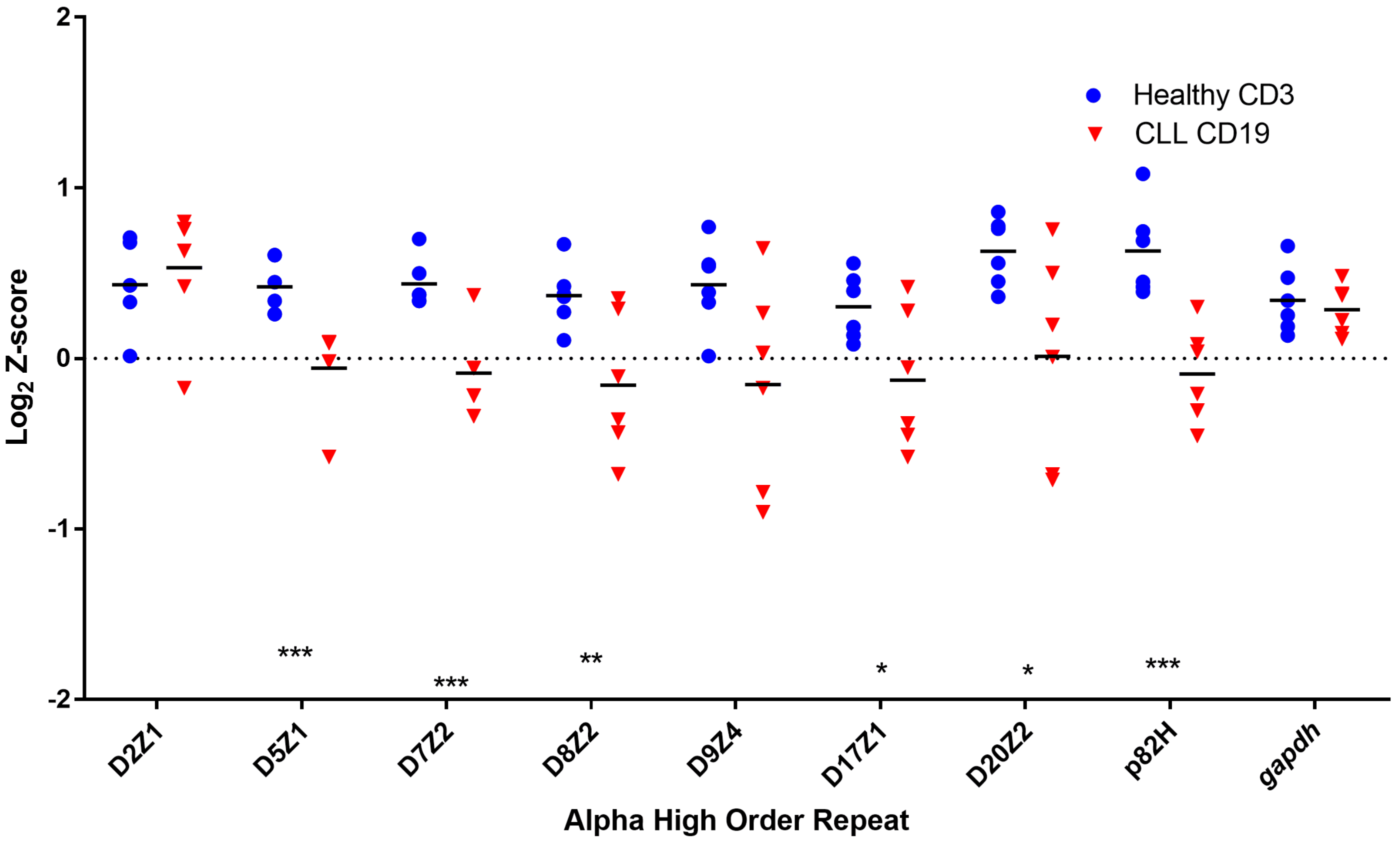


**Supplementary Figure S6.** Heterogeneous loss of centromeres in CLL patients. Dot plot representing the abundance of α-satellites specific for each centromere array (X axis) in CLL tumors. Abundance of centromere specific α-satellites is depicted by the $\log_2$ Z-score (Y-axis) of each α-satellite. The $\log_2$ normalized numbers for each α-satellite were normalized to the average copy number of a given repeat in DNA from healthy cells (blue circles). Statistical significance was calculated using a t-test. * = $p < 0.05$, ** = $p < 0.01$, *** = $p < 0.001$, **** = $p < 0.0001$.
